# On a ‘failed’ attempt to manipulate conscious perception with transcranial magnetic stimulation to prefrontal cortex

**DOI:** 10.1101/198218

**Authors:** Eugene Ruby, Brian Maniscalco, Hakwan Lau, Megan A.K. Peters

**Author notes:** **Correspondence should be addressed to:** Eugene Ruby UCLA Psychology Department 1285 Franz Hall, Box 951563 Los Angeles, CA 90095-1563.

## Abstract

It has been reported that continuous theta burst transcranial magnetic stimulation (TMS) to the dorsolateral prefrontal cortex (DLPFC) impairs metacognitive awareness in visual perception (Rounis et al., 2010). Bor et al. (2017) recently attempted to replicate this result. However, the authors modified the experimental design of the original study considerably, meaning that this was not strictly a replication. In some cases, the changes are *a priori* expected to lower the chance of obtaining positive findings. Despite these changes, the researchers in fact still found an effect similar to Rounis et al.’s, but they claimed that it was necessary to adopt certain criteria to discard ∽30% of their subjects, after which a null result was reported. Using computer simulations, we evaluated whether the subject exclusion criteria Bor et al. adopted was appropriate or beneficial. We found that, contrary to their intended purpose, excluding subjects by their criteria does not actually reduce false positive rates. Taking into account both their positive result (without subject exclusion) and negative result (after exclusion) in a Bayesian framework, we further found that their results suggest a 75% or greater likelihood that TMS to DLPFC does in fact impair metacognition, directly contradicting their claim of replication failure. That lesion and chemical inactivation studies are known to demonstrate positive effects in similar paradigms further suggests that Bor et al.’s alleged negative finding cannot be taken as evidence against the role of the prefrontal cortex in conscious perception in general.

## Introduction

Visual metacognition refers to how well one can give subjective judgments to discriminate between correct and incorrect perceptual decisions (Fleming & Lau, 2014). As visual metacognition appears to be closely linked to conscious awareness (Ko & Lau, 2012, Maniscalco & Lau, 2016), it is of interest that the prefrontal cortex has been heavily implicated in mediating both of these faculties (Baird Smallwood, Gorgolewski, & Margulies, 2013; Del Cul, Dehaene, Reyes, Bravo, & Slachevsky, 2009; Fleming, Ryu, Golfinos, & Blackmon, 2014; Lau & Passingham, 2006; Lumer, Friston, & Rees, 2008; McCurdy et al., 2013; Rounis, Maniscalco, Rothwell, Passingham, & Lau, 2010; Turatto, Sandrini, & Miniussi, 2004). Specifically, one prefrontal area with an empirical link to these abilities is the dorsolateral prefrontal cortex (DLPFC; Lau and Passingham, 2006; Rounis et al., 2010, Turatto et al., 2004).

It has been reported that continuous theta-burst transcranial magnetic stimulation (TMS) to DLPFC can impair visual metacognition (Rounis et al., 2010). Recently, Bor, Schwartzman, Barrett, & Seth (2017) attempted to replicate this finding but reported a null result, which they took to suggest that DLPFC might not be “critical in generating conscious contents” (Bor et al., 2017, p. 16).

However, although the researchers motivated their experiments as direct replications (e.g., Bor et al., 2017, p. 3), they made several changes to the original study design, some of which are known to undermine the chance of finding meaningful results from the outset. In particular, in their main experiment (their Experiment 1) the researchers used a between-subjects design instead of the within-subjects design used by Rounis et al. (2010), which might have limited their statistical power (Greenwald, 1976). Although they attempted to address this potential issue in a second study (their Experiment 2), we will show below that this study design was unsatisfactory for other reasons.

Importantly, despite these modifications, Bor et al. (2017) in fact found a positive result akin to Rounis et al.’s (2010), with both studies reporting comparable changes in metacognition for subjects who received TMS to DLPFC. However, the researchers proceeded to set stringent exclusion criteria, which they claimed should lower false positive rates (i.e., rate of incorrectly detecting an effect). This caused the removal of a relatively large number of subjects (27 out of 90), and ultimately a null result was found, leading Bor et al. to conclude that the initial significant finding must have been spurious. But the exact criteria for subject exclusion seem arbitrary; therefore, here we formally evaluate the consequences of adopting such criteria in a simulation, and what the interpretation of their results should have been in a Bayesian framework.

## Methods

Our goal was to assess whether excluding subjects as in Bor et al. (2017) was needed to curb false positive rates as they claim, and also whether doing so led to increased false negative rates and thus lower statistical power (i.e., probability of successfully detecting a true effect). Therefore, we simulated two populations of subjects, one that exhibited the TMS-induced metacognitive impairments, as in Rounis et al. (2010), and one that did not, and included them in two sets of 1,000 “experiments” that mirrored Bor et al.’s Experiment 1 (between-subjects design). We then compared statistical power and false positive rates both before and after implementing the exclusion criteria used by Bor et al.

### Building two populations of “subjects”

Each simulated “subject” was characterized by four parameters to produce behavioral outputs with and without TMS. For the first three parameters -- objective performance capacity (d’*_s_*), response bias (Type 1 criterion; *c*_*s,1*_), and subjective response biases (Type 2 criteria; *c*_*s,2*_) -- the values were taken from Rounis et al. (2010) to mimic the distributions seen there. These values were then fixed for each simulated subject across task conditions (pre- and post-TMS).

To simulate the effect of degraded metacognitive sensitivity by TMS, we defined a fourth parameter corresponding to Type 2 (i.e. metacognitive) noise (σ*_s,TMS_*), such that in the TMS condition Type 2 criteria (*c*_*s,2*_) become unstable and the trial-by trial correspondence between confidence and accuracy is lowered (Maniscalco & Lau, 2012; Maniscalco & Lau, 2016; Peters et al., 2017). Thus, for each simulated subject for these TMS conditions, over trials we added TMS noise (σ*_s,TMS_*) to the subject’s internal response, after their discrimination judgment but before their confidence judgment (see task description in Figure 1), such that across all simulated subjects the average reduction in metacognitive sensitivity mimicked that found in Rounis et al. (2010; see supplemental materials for more details).

**Figure 1.**
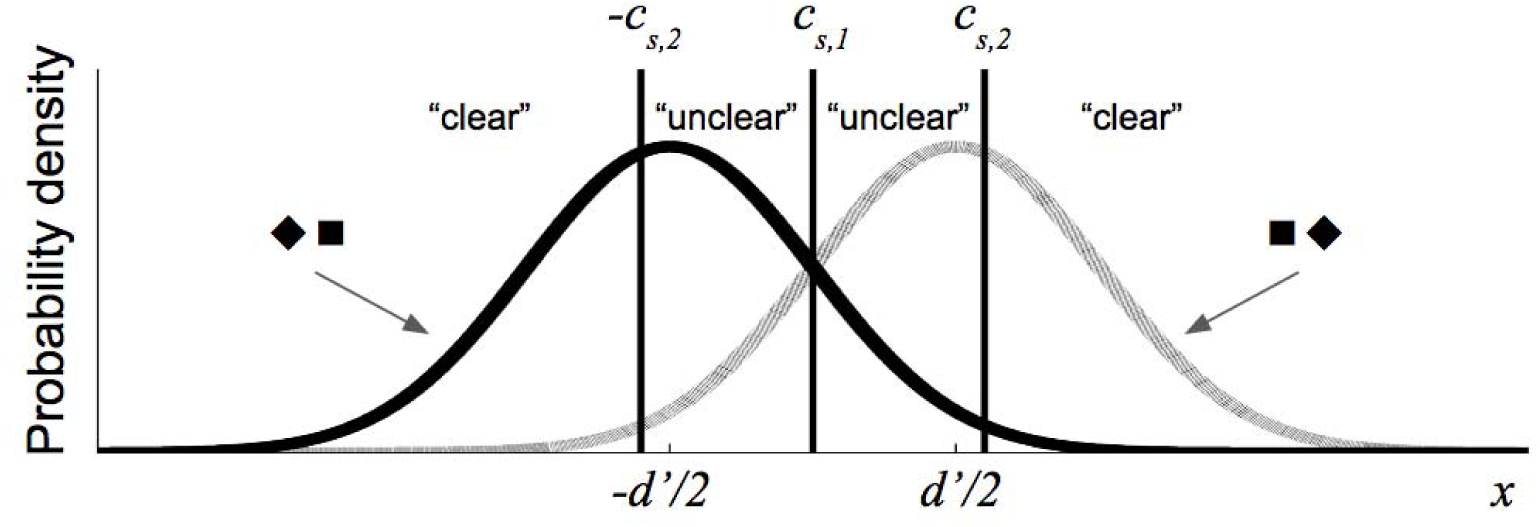
Signal detection theoretic framework for the simulated spatial 2AFC task. For a given subject s, each stimulus presentation (either ▪ ♦ or ♦▪) caused an internal reponse value (x), with X _▪ ♦_∽ N(d’/2,1) and X _♦▪_∽ N(-d’/2,1) and the subject then indicated which of the two shape configurations appeared. If x exceeds the subjects Type 1 criterion (c_s_, 1), then the subject responded “▪ ♦;” otherwise the subject responded “♦▪.” Objective performance capacity (d’) is the normalized distance between the two distributions. The subject then indicated how clearly / confidently they saw the stimulus, based on a comparison between x and the Type 2 criterion (-c_s,2_ and c_s,2_). If x < c_s,1_ or x > c_s,2_ the subject responded “clear;” otherwise the subject responded “unclear.” If TMS is present and assumed to degrade metacognitive sensitivity, noise(σ _s,TMS_) is added for the Type 2 responses, such that the relevant computations are whether x < c_s,1_ + ɛ_trial_ or x > c_s,2_ + ɛ_trial_, with ɛ_trial_ ∽ N(0, σ_s,TMS_) and resampled on each trial (see supplemental materials for more details).

### Simulating the behavioral task

Our simulated task followed the spatial two-alternative forced-choice (2AFC) task design used by Rounis et al. (2010) and Bor et al. (2017) (see Figure 1).

### Assessing Metacognitive Performance

For all simulated subjects, we calculated meta d’, a bias-free measure of metacognitive sensitivity (Maniscalco & Lau, 2012), using a standard toolbox (Maniscalco, 2014) whereby estimation was conducted by minimizing the sum of squared errors (SSE), as in Rounis et al. (2010) and Bor et al. (2017). As with both of these studies, our measure of interest was meta d’-d,’ which indicates a participant’s metacognitive sensitivity for a given level of basic task performance (Fleming and Lau, 2014).

### Simulating populations of subjects

Before running our simulated “experiments,” we first built two populations of subjects: one designed to show the impairing effect of TMS on metacognitive performance (“effect-present population”), based on the results found in Rounis et al. (2010), and the other showing no such effect (“effect-absent population”; such that the above mentioned parameter σ*_s,TMS_* was set to 0). The two populations each contained 10,000 subjects (5,000 completing the Pre-Real-TMS and Post-Real-TMS conditions and 5,000 completing the Pre-Sham-TMS and Post-Sham-TMS conditions). As described above, between-subjects variability in the simulation parameters was based on the empirical between-subject variability reported by Rounis et al. (2010).

To verify that the effect on metacognitive sensitivity reported by Rounis et al. (2010) was successfully recreated in our simulated effect-present population and not in the effect-absent population, we compared the means for each condition and also ran a mixed-design ANOVA on metacognitive sensitivity (meta d’ - d) for each population, with between-subjects factor TMS type (Real TMS, Sham TMS) and within-subject factor time (Pre-TMS, Post-TMS). We confirmed the impairing effect of TMS on metacognitive sensitivity in the effect-present population with a significant TMS type x time interaction; F(1,9998) = 307.86, p < .001, and the means for each condition were as follows: for group 1, Pre-Real-TMS d’ = 1.698 and meta d’ = 1.557, Post-Real-TMS d’ = 1.713 and meta d’ = 1.172; for group 2, Pre-Sham-TMS d’ = 1.678 and meta d’ = 1.540, Post-Sham-TMS d’ = 1.698 and meta d’ = 1.577. Conversely, we confirmed no impairing effect of TMS on metacognitive sensitivity for the effect-absent population; F(1,9998) = 1.79, p = 0.18, and condition means were as follows: for group 1, Pre-Real-TMS d’ = 1.680 and meta d’ = 1.576, Post-Real-TMS d’ = 1.694 and meta d’ = 1.580; for group 2, Pre-Sham-TMS d’ = 1.685 and meta d’ = 1.534, Post-Sham-TMS d’ = 1.702 and meta d’ = 1.565.

### Simulating Bor et al.’s (2017) Experiment 1

As in Bor et al.’s (2017) between-subjects experiment, the simulated subjects were randomly assigned to either the Real or Sham TMS group. As in their study, each condition contained 300 trials of the spatial 2AFC task.

We simulated 1,000 ‘experiments’, each containing samples of 35 subjects drawn with replacement from both the corresponding effect-present and effect-absent populations. Subjects in each sample were randomly assigned to one of two groups: 17 subjects were exposed to the two real TMS conditions and 18 subjects were exposed to the two sham TMS conditions, as in Bor et al.’s (2017).

For each simulated experiment, we first performed statistical tests with all subjects. Each ‘experiment’ was analyzed with a mixed-design ANOVA on metacognitive sensitivity (meta d’-d’) with between-subjects factor TMS type (Real TMS, Sham TMS) and within-subject factor time (Pre-TMS, Post-TMS); a ‘positive’ effect of TMS on metacognitive sensitivity was found if the interaction between TMS type and time was found to be significant (p<.05).

We then performed the same tests after excluding subjects using Bor et al’s criteria (Type 1 and/or Type 2 hit rates or false alarm rates < 0.05 or > 0.95, or with Type 1 percent correct values less than or equal to 65%). Similarly to Bor et al. (2017), we excluded about 30% of subjects on average for all 1,000 simulations for both our effect-present simulations (mean = 30.20%, S.D. = 7.77%) and effect-absent simulations (mean = 30.86%, S.D. = 7.96%).

We assessed the consequences of excluding subjects on statistical power by examining the percentage of simulated experiments done on the effect-present population for which the TMS x time interaction correctly reached significance (p < .05) when subjects were not excluded (as done by Rounis et al., 2010) versus excluded (as done by Bor et al., 2017).

(We also simulated other experiments, and the details are in Supplementary Materials.)

## Results

The results of simulations based on Bor et al.’s (2017) Experiment 1 showed that false positive rates were nearly identical for non-exclusion and exclusion (0.046 and 0.048, respectively; Figure 2a, Table 1). This suggests that excluding subjects has no effect on false positive rates, contrary to what Bor et al. (2017) claimed.

**Figure 2.**
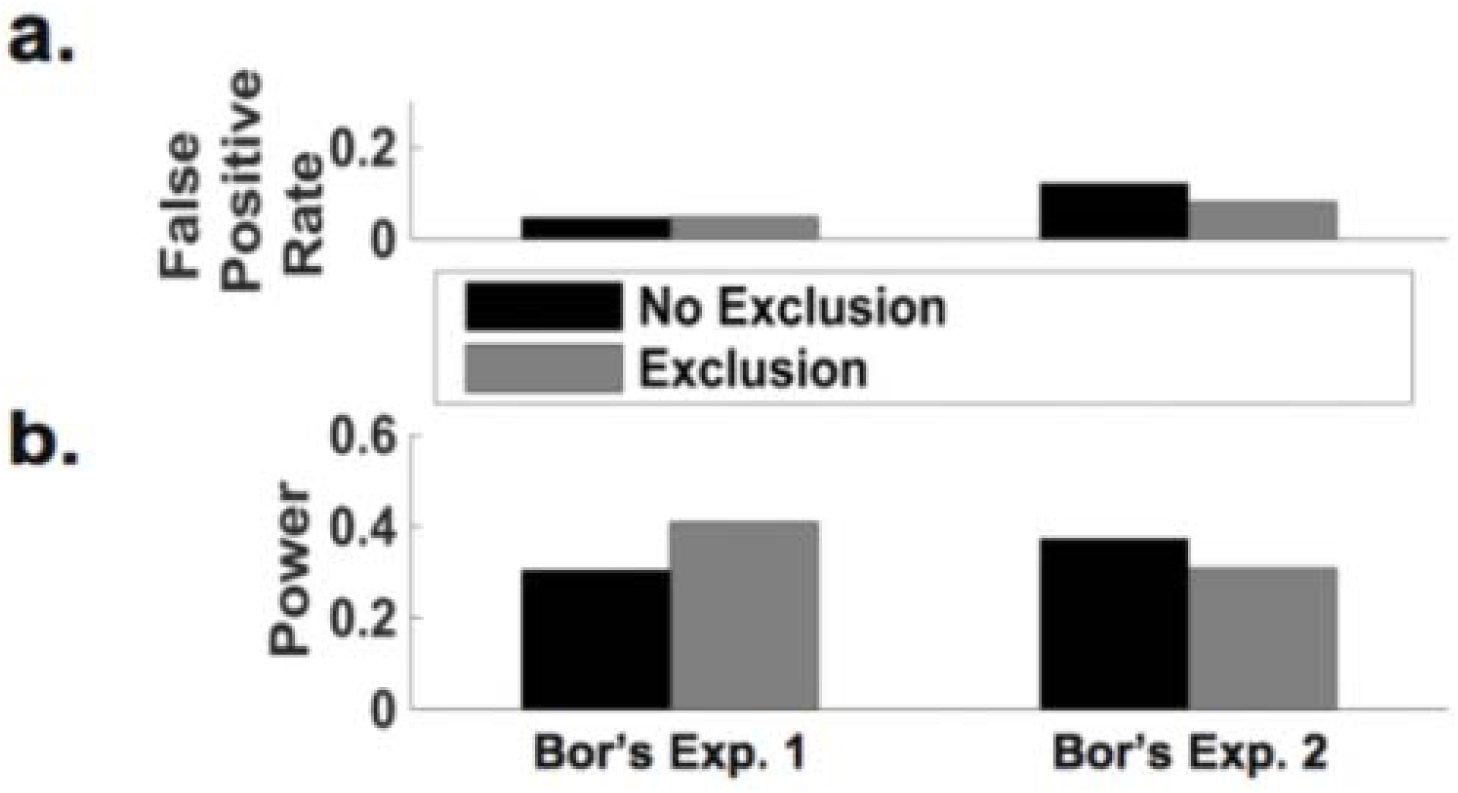
Excluding subjects yields little impact on statistical power or false positive rates. As in the actual studies, the effect of TMS was assessed by the statistical significance of the interaction between TMS (DLPFC or control) and time (before TMS and after TMS). (a) Excluding versus mot excluding subjects yielded no meaningful change in false positive rate for the between – subjects design (Bor’s Exp. 1). Exclusion led to a small decrease in false positive rate for the double – repeat design (Bor’s Exp 2), but this design inflated false positive rate in comparison to the between – subjects design (Bor Exp. 1). (b) Compared with not excluding subjects, excluding subjects yielded an increase in statistical power for the between – subjects design (Bor’s Exp. 1). Conversely, exclusion resulted in a decrease in power in comparison with no exclusion for the unconventional double – repeat within – subjects design (Bor’s Exp. 2). Importantly, the double – repeat design (Bor’s Exp. 2) led to considerably lower rather than the intended higher statistical power.

Interestingly, excluding subjects led to an increase in power (power_no exclusion_ = 0.304 vs. power_exclusion_ = 0.409; Figure 2b, Table 1), contrary to what might be expected from a reduced sample size. However, we note that despite this modest increase, power is still fairly low; given if an effect is present one is more likely to miss it than to not miss it.

### What should be the correct interpretation given the results?

We used Bayesian analysis to calculate the probability that TMS actually caused metacognitive sensitivity deficits in the population of human subjects tested by Bor et al. (2017), given the pattern of results they observed. We used Bayes’ rule,

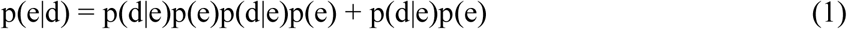

where the posterior probability *p*(*e|d*) refers to the probability of a true effect *e* on metacognitive sensitivity due to TMS given the observed data *d*, as a function of the likelihood of observing the present data given the effect is true, *p*(*d|e*), and the prior probability of the effect being true, *p*(*e*). We assign the observed data *d* as being a significant interaction *before* excluding subjects that then disappeared when subjects were excluded, as reported by Bor et al. (2017). So as to not bias the calculation in either direction, we assign equal prior probability (0.5) to the effect being present or not at the population level.

In 8.9% of simulated experiments, an effect was correctly detected when no subjects were excluded, but subsequently excluding subjects led to a false negative (i.e., *p*(*d|e*)). In 2.9% of simulated experiments a false positive was initially found when not excluding subjects, but subsequently excluding subjects resulted in correctly finding no effect (i.e., *p*(*d|∽e*)). Putting these values into Bayes’ rule (Equation 1), we find that the probability of TMS actually causing metacognitive deficits at the population level given that an effect was observed but then disappeared to be high, *p*(*e|d*) = .7542. This is because the probability that an effect was observed (without exclusion) but then disappeared with exclusion is 3.07 times higher when the effect was actually present in the population than when it was absent.

In other words, had one not been initially persuaded by Rounis et al (2010)’s findings, and assumed that there was only half a chance of the effect’s being true (0.5), upon seeing Bor et al’s (2017) pattern of results one should update this belief -- if one were rational -- to recognize that the effect is 75.42% likely to be true. This is exactly the opposite of what Bor et al. concluded. Moreover, we should be even surer of this conclusion if we take Rounis et al.’s findings at face value. For example, setting the prior at .95 would yield a 98.31% posterior probability of the effect being true.

### Simulating Other Experiments

Bor et al. (2017) acknowledged that their Experiment 1 may have lacked power due to using a between-subjects instead of a within-subjects design (despite our findings above); therefore, they ran a second experiment using an unusual “double-repeat within-subjects” design. This design involved up to four days of real and sham TMS manipulations (two days of each) in which subjects only advanced to subsequent days if their performance met certain benchmarks. To assess whether this unconventional design actually increased power, we ran a second simulation akin to what we did above (described in detail in supplementary materials).

Contrary to what Bor et al. (2017) intended, following their subject exclusion procedure we actually observed a *decrease* in power from 0.409 in the between-subjects design to 0.308 in their double-repeat design (Figure 2b). Also, there was an increase in the false positive rate from 0.048 (in the between-subjects design) to 0.081 (in the double-repeat design). Moreover, for this unconventional double-repeat design we found that not excluding subjects actually yielded higher power (0.372) than excluding subjects (0.308; Figure 2b). Also, false positive rates were slightly higher for exclusion (0.081) than for non-exclusion (0.121; Figure 2a).

In fact, had Bor et al. (2017) simply used the same sample size in this second experiment (n = 27) to actually replicate Rounis et al.’s (2010) simple within-subject experiment, the authors would have achieved their intended purpose. We ran an additional simulation and showed that implementing this design with n = 27 would have increased power to 0.567. We found similar false positive rates for exclusion and non-exclusion (0.053 and 0.053, respectively), although as in Bor et al.’s Experiment 1 power was higher for exclusion (0.567) in comparison with non-exclusion (0.388).

For completeness we ran a final simulation to assess whether exclusion could have been useful in Rounis et al.’s (2010) original study with original sample size of n = 20. We found similar false positive rates for exclusion and non-exclusion (0.048 and 0.042, respectively), and that power was higher for exclusion than non-exclusion (0.432 and 0.311, respectively).

(See Table 1 in supplementary materials for a complete listing of results for all experimental designs.)

## Discussion

We found that Bor et al. (2017) did not achieve their main goal of causing meaningful reductions in statistical false positives by the use of their subject exclusion criteria. Such exclusion is not necessary because false positive rates were low to begin with, reflecting the general robustness of this kind of statistical analysis. Although in some instances excluding subjects improved power, presumably by removing noisy outliers, the resulting power is still low and this improvement does not always happen (e.g. the double repeat-design). Most importantly, upon seeing the pattern of results in Bor et al’s Experiment 1 (positive result turned negative after exclusion), we showed with Bayesian analysis that the correct interpretation should be to conclude that the effect is very likely to be present, contrary to their claims.

While Bor et al. (2017) acknowledge that their Experiment 1 may lack power, because it uses a between-subject design, their attempt to alleviate this with their Experiment 2 (an unconventional ‘double-repeat’ design) did not work well. In particular, after using their exclusion criterion, power actually decreased in that experiment, and false positive rate also increased slightly.

The supposed goal of Bor et al.’s (2017) series of experiments was to directly replicate Rounis et al. (2010). We think such effort is important and should be applauded, and in fact Rounis et al. shared their source code with Bor et al. in support of their endeavor. However, Bor et al. puzzlingly changed several elements of the original study. For example, they used confidence ratings instead of the visibility ratings used by Rounis et al. It’s possible that the effect of TMS to prefrontal cortex may work better with visibility rather than confidence judgments, as it is known that different subjective measures can lead to systematically different results (Sandberg, Timmermans, Overgaard, & Cleeremans, 2010; King & Dehaene, 2014). Specifically in the context of metacontrast masking, Maniscalco & Lau (2016) recently replicated a dissociation between stimulus discrimination performance and visual awareness (Lau & Passingham, 2006) when using visibility ratings, but not when utilizing confidence ratings (personal communication). Another notable change is that Bor et al. didn’t instruct subjects to use their two metacognitive ratings evenly -- unlike Rounis et al., who did give this instruction.

If Bor et al. (2017) were concerned about the robustness of meta d’ estimations, they could have used more than two levels of metacognitive ratings, which is commonly done in the literature (Overgaard, 2015). Using only two metacognitive rating levels (as both studies did) will only provide one point on the Type 2 receiver operating characteristic (ROC) curve, which may limit the efficiency of meta d’ estimations (Maniscalco & Lau, 2012). Using more than two rating levels helps because even if one (or even two) point on the Type 2 ROC curve is extreme, it is unlikely that the other point(s) will also be so extreme. This point should be easily appreciated by Bor et al., as one of the authors’ own analysis (Seth, 2011) showed that they would have had to use many more trials (i.e., thousands) to accurately estimate of meta d’ if there were only two rating levels.

These considerations would probably have made the impression of needing to exclude subjects unnecessary -- which we now show is the case regardless. From the outset, it may seem correct to exclude subjects on the basis that including the “unstable” data violated the assumption of normality in Bor et al.’s (2017) between-subjects experiment. But ultimately, assumptions in statistical inferences are never meant to be perfectly realistic and precise. What matters is whether the corresponding inference may be correct or not based on the data. Our analysis shows that subject exclusion serves no benefit in improving the validity of such inferences when it comes to false positive rates, contrary to what Bor et al. intended. Further supporting this point, we used the Shapiro-Wilk test on distributions of metacognitive sensitivity (meta d’ - d’) for the 1000 effect-absent simulations and found that significant violations of normality were more prevalent without subject exclusion (95.6% of simulations) than with exclusion (73.0% of simulations). Thus, although subject exclusion tends to produce distributions that are more Gaussian (consistent with Bor et al.’s findings), the improvement in normality due to exclusion was not accompanied by a reduction in false positive rates. These simulation findings are inconsistent with Bor et al.’s conclusion that the significant effect found in their data prior to subject exclusion was a false positive driven by non-normality of the data.

We recognize that TMS is limited in sensitivity compared to more invasive methods. In fact, power in the concerned studies is in general quite low according to our simulations. Also, neuronavigation techniques (e.g., Brainsight TMS Navigation) are probably needed to ensure targeting precision of stimulation -- a technique that neither Rounis et al (2010) nor Bor et al. (2017) used. Of relevance, Rahnev, Nee, Riddle, Larson, & D’Esposito (2016) recently reported that theta-burst TMS to DLPFC actually increased metacognitive performance, which the authors suggest may have been due to the fact that a very anterior portion of DLPFC was stimulated for most study participants. This may suggest that different parts of DLPFC perform different functions.

Taken together, these considerations suggest that replicating the findings of Rounis et al (2010) is possibly non-trivial. It is therefore striking that Bor et al (2017) in fact succeeded in doing so, despite their interpreting otherwise, and despite the various changes they have made to the design of the experiment including using a between-subject design with limited power.

Ultimately, whether theta-burst TMS to DLPFC can robustly impair visual awareness concerns the specific method and details. More important is the general question regarding the role of the prefrontal cortex in visual awareness (Odegaard, Knight, & Lau, in press; Lau & Rosenthal, 2011). Bor et al. (2017) claim that lesions to the prefrontal-parietal network (PPN) tend to show “at best subtle impairments in conscious detection” (Del Cul et al., 2009; Simons, Peers, Mazuz, Berryhill, & Olson, 2010). However both of these cited studies actually found positive results, and it is unclear by what standard Bor et al. (2017) judge them to be “subtle.” Contradictorily, in a 2012 review, Bor & Seth (2012) themselves cite Del Cul et al. (2009) and Simon et al. (2010), among other positive results, claiming that these findings “strongly implicate all key individual components of the PPN in conscious processing” (Bor & Seth, 2012, p. 3).

Importantly, in another study adopting similar psychophysical measures and task procedures as Bor et al. (2017), Fleming et al. (2014) found a ∽50% decrease in metacognitive efficiency for patients with prefrontal lesions. Supposedly, a 50% decrease should not be considered a small, subtle, effect. Furthermore, Cortese, Amano, Koizumi, Kawato, & Lau (2016) showed that manipulation of PFC activity via biofeedback of decoded fMRI information can alter metacognitive confidence ratings. Also, chemical inactivation of the PFC has recently been shown to induce deficits in metamemory in monkeys (Miyamoto et al., 2017).

While we think the current attention to the issue of replicability within psychology and cognitive neuroscience is a most useful and important development, we worry that this may occasionally generate undue excitement in some apparent non-replications. We hope that the above discussion makes clear that the alleged non-replication by Bor et al. (2017), even if true (which we have shown is unlikely), does not meaningfully speak to the role of PFC in conscious perception in general.

## References

Baird, B., Smallwood, J., Gorgolewski, K. J., & Margulies, D. S. (2013). Medial and Lateral Networks in Anterior Prefrontal Cortex Support Metacognitive Ability for Memory and Perception. J Neurosci, 33(42), 16657–65.

Barrett, A., Dienes Z., & Seth, A. K. (2013). Measures of metacognition on signal-detection theoretic models. Psychol Methods, 18(4): 535–52.

Bor, D. & Seth, A. K. (2012). Consciousness and the Prefrontal Parietal Network: Insights from Attention, Working Memory, and Chunking. Front Psychol, 3: 63.

Bor, D., Schwartzman, D. J., Barrett, A. B., & Seth, A. K. (2017). Theta-burst transcranial magnetic stimulation to the prefrontal or parietal cortex does not impair metacognitive visual awareness. PLoS One, 12(2): e0171793.

Cortese, A., Amano, K., Koizumi, A., Kawato, M., & Lau, H. (2016). Multivoxel neurofeedback selectively modulates confidence without changing perceptual performance. Nat Commun, 7: 13669.

Del Cul, A., Dehaene, S., Reyes, P., Bravo, E., & Slachevsky, A. (2009). Causal role of prefrontal cortex in the threshold for access to consciousness. Brain, 132(Pt 9): 2531–40.

Greenwald, A. G. (1976). Within-subjects designs: to use or not to use? Psychol Bull, 83(2): 314–20.

Fleming, S. M. & Lau, H. (2014). How to measure metacognition. Front Hum Neurosci, 8: 443.

Fleming, S. M. Ryu, J. Golfinos, J. G. & Blackmon, K. E. (2014). Domain-specific impairment in metacognitive accuracy following anterior prefrontal lesions. Brain, 137(Pt 10): 2811–22.

King, J. R. & Dehaene, S. (2014). A model of subjective report and objective discrimination as categorical decisions in a vast representational space. Philos Trans R Soc Lond B Biol Sci., 369(1641): 20130204.

Ko, Y. & Lau, H. (2012). A detection theoretic explanation of blindsight suggests a link between conscious perception and metacognition. Philos Trans R Soc Lond B Biol Sci., 367(1594): 1401–11.

Lau, H. C. & Passingham, R. E. (2006). Relative blindsight in normal observers and the neural correlate of visual consciousness. Proc Natl Acad Sci, USA, 103(49):18763–8.

Lau, H. & Rosenthal, D. (2011). Empirical support for higher-order theories of conscious awareness. Trends Cogn Sci, 15(8): 365–373.

Lumer, E. D., Friston, K. J., & Rees, G (1998). Neural correlates of perceptual rivalry in the human brain. Science, 280(5371): 1930–4.

Macmillan, N. A. & Creelman, C. D. (2004). Detection theory: a user’s guide. Mahwah, NJ: Lawrence Erlbaum.

Maniscalco, B. & Lau, H. (2012). A signal detection theoretic approach for estimating metacognitive sensitivity from confidence ratings. Conscious Cogn., 21(1): 422–30.

Maniscalco, B (2014, October). Type 2 signal detection theory analysis using meta-d’. Retrieved from http://www.columbia.edu/∽bsm2105/type2sdt/

Maniscalco, B. & Lau, H. (2016). The signal processing architecture underlying subjective reports of sensory awareness. Neurosci Conscious, 2016(1): niw002.

McCurdy, L. Y., Maniscalco, B., Metcalfe, J., Liu, K. Y., de Lange, F. P., & Lau, H. (2013). Anatomical Coupling between Distinct Metacognitive Systems for Memory and Visual Perception. J Neurosci, 33(5): 1897–906.

Miyamoto, K., Takahiro, O., Setsuie, R., Takeda, M., Tamura, K., Adachi, Y., Miyashita, Y. (2017). Causal neural network of metamemory for retrospection in primates. Science, 355(6321): 188–193.

Odegaard, B., Knight, R., & Lau, H. (in press). Should A Few Null Findings Falsify Prefrontal Theories of Consciousness? J Neurosci.

Overgaard, M. (2015). Behavioral Methods in Consciousness Research. Oxford, United Kingdom: Oxford University Press. Print.

Peters, M. A. K., Thesen, T., Ko, Y. D., Maniscalco, B., Carlson, C., Davidson, M., Doyle, W., Kuzniecky, R., Devinsky, O., Halgren, E., & Lau, H. (2017). Perceptual confidence neglects decision-incongruent evidence in the brain. Nat Hum Behav, 1: 0139.

Rahnev, D., Nee, D. E., Riddle, J., Larson, A. S. & D’Esposito, M. (2016). Causal evidence for frontal cortex organization for perceptual decision making. Proc Natl Acad Sci U S A, 113(2): 6059–6064.

Rounis, E., Maniscalco, B., Rothwell, J. C., Passingham, R. E., & Lau, H. (2010). Theta-burst transcranial magnetic stimulation to the prefrontal cortex impairs metacognitive visual awareness. Cogn Neurosci, 1(3): 165–75.

Sandberg, K., Timmermans, Overgaard M., Cleermans, A. (2010). Measuring consciousness: Is one measure better than the other? Conscious Cogn, 19(4): 1069–78.

Seth, A. K. (2011, June). Signal detection theory is not a good model or measure of consciousness. Presented at the 15th Annual Meeting of the Association for the Scientific Study of Consciousness, Kyoto, Japan.

Simons, J. S., Peers, P. V., Mazuz, Y. S., Berryhill, M. E., & Olson, I. R. (2010). Dissociation between memory accuracy and memory confidence following bilateral parietal lesions. Cereb Cortex, 20(2): 479–85.

Turatto M., Sandrini M., Miniussi C. (2004). The role of the right dorsolateral prefrontal cortex in visual change awareness. Neuroreport, 15(16): 2549–52.

